# Microscale bulk segregant analysis

**DOI:** 10.64898/2026.10.09.757075

**Authors:** Giancarlo N. Bruni, Lauren C. Crisman, Leslie Alamo, Heriberto Marquez, James Boocock, Joshua S. Bloom, Leonid Kruglyak

**Author notes:** Correspondence (GNB), (JSB), (LK).

## Abstract

The traits of an organism depend on its genetic background and environment. Bulk segregant analysis (BSA) maps quantitative trait loci (QTL) by measuring parental allele frequencies in pools of genetically recombined progeny (segregants). Conventional BSA requires laborious separate sample handling for each cross and environmental condition. Here, we describe a 96-well liquid format for extreme QTL mapping (X-QTL) in *Saccharomyces cerevisiae*. We crossed 40 isolates to a common parent carrying mating-type-specific drug-resistance markers at *HO*, which enabled us to select diploids and haploid segregants in 1 mL cultures. Each well held one segregant pool, which we then grew in deep wells in four conditions: the standard glucose medium, sodium chloride or caffeine added to the standard medium, or acetate replacing glucose as a carbon source. We barcoded DNA samples from each of 384 wells and sequenced them in one Illumina Nova-seq X run. Sequencing data for 181 segregant pools passed quality control steps and was used to map 1,472 QTL in 35 crosses. We replicated a subset of QTL previously mapped in these growth conditions by conventional X-QTL. The 96-well approach increases the throughput of every step of X-QTL, enabling genetic analysis of many more cross and conditions.

**ARTICLE SUMMARY:** The traits of an organism are shaped by its genes and environment. A standard approach to find the genetic differences that underlie trait variation in model organisms is to mate two genetically distinct individuals, grow a large number of their offspring in an environmental condition of interest, and compare the frequencies of parental alleles across the genome between a control pool of offspring and a pool that represents a phenotypic tail of improved growth in that condition. We developed a method to adapt this approach to small liquid volumes, enabling genetic analysis of many crosses and conditions in parallel. We applied the method to 40 crosses and 4 conditions and found many regions of the genome where a difference between the parents influenced growth.

## INTRODUCTION

QTL mapping in *Saccharomyces cerevisiae* has historically used two approaches. One uses panels of individually genotyped and phenotyped segregants from one cross (Brem *et al*. 2002; Steinmetz *et al*. 2002; Deutschbauer and Davis 2005; Perlstein *et al*. 2007; Bloom *et al*. 2013, 2015) or several crosses (Cubillos *et al*. 2011; Bloom *et al*. 2019). These panels give direct estimates of heritability, effect sizes and gene–gene interactions, but every segregant must be isolated, genotyped and phenotyped. In the second approach, bulk segregant analysis (BSA), segregants are instead pooled, the segregant pool is selected for a phenotypic tail of interest, and QTL are detected as differences in allele frequency between selected and unselected control segregant pools genotyped in bulk (Michelmore *et al*. 1991; Brauer *et al*. 2006; Segrè *et al*. 2006). We previously developed extreme QTL mapping (X-QTL), which applies BSA to pools of millions of segregants (Ehrenreich *et al*. 2010), and used it to map a number of quantitative traits (Ehrenreich *et al*. 2010, 2012; Torabi and Kruglyak 2012; Albert *et al*. 2014; Treusch *et al*. 2015).

Pooled approaches have been used to map many traits in two-parent crosses (Birkeland *et al*. 2010; Wenger *et al*. 2010; Swinnen *et al*. 2012; Hubmann *et al*. 2013; Pais *et al*. 2013; Yang *et al*. 2013; Duitama *et al*. 2014; Wilkening *et al*. 2014; Hou *et al*. 2016; Meijnen *et al*. 2016; Fernández-Niño *et al*. 2018; Higgins *et al*. 2018; She and Jarosz 2018; Sirr *et al*. 2018; Brion *et al*. 2020; Stojiljkovic *et al*. 2020; Coradini *et al*. 2021). They have also been applied to advanced-intercross, multiparent and hybrid populations (Parts *et al*. 2011; Cubillos *et al*. 2013, 2017; García-Ríos *et al*. 2017; Barré *et al*. 2020; Naseeb *et al*. 2021; Visinoni *et al*. 2024), as well as combined with backcrossing (Taylor and Ehrenreich 2014; Matsui *et al*. 2015; Taylor *et al*. 2016; de Bem *et al*. 2026) or with fixation of one chromosome (Buzby *et al*. 2025). In all of these studies, every cross in every condition was grown and analyzed separately, and thus the overall effort scales with the number of crosses and conditions.

Here, we describe a 96-well liquid format for X-QTL. We perform mating, sporulation, random-spore preparation, selection, DNA extraction, and Tn5 library construction in plates, so that each well is a self-contained experiment of a segregant pool in a condition of interest. One 96-well plate holds 96 such segregant pools, greatly increasing scalability. We mapped QTL in crosses between 36 isolates and one shared parent strain in four conditions and detected previously reported and new QTL.

## RESULTS

### A 96-well liquid format for X-QTL

X-QTL has three major steps: a large pool of segregants is generated from a cross between two strains, the pool is selected for extreme phenotypes, and allele frequencies across the genome are measured in the selected pool and in an unselected control pool (Ehrenreich *et al*. 2010) (Figure 1). We carried out every X-QTL step in 96-well plates, with each well holding a pool of segregants from one cross growing in one condition. Three modifications of the protocol allowed for this miniaturization.

**FIGURE 1.**
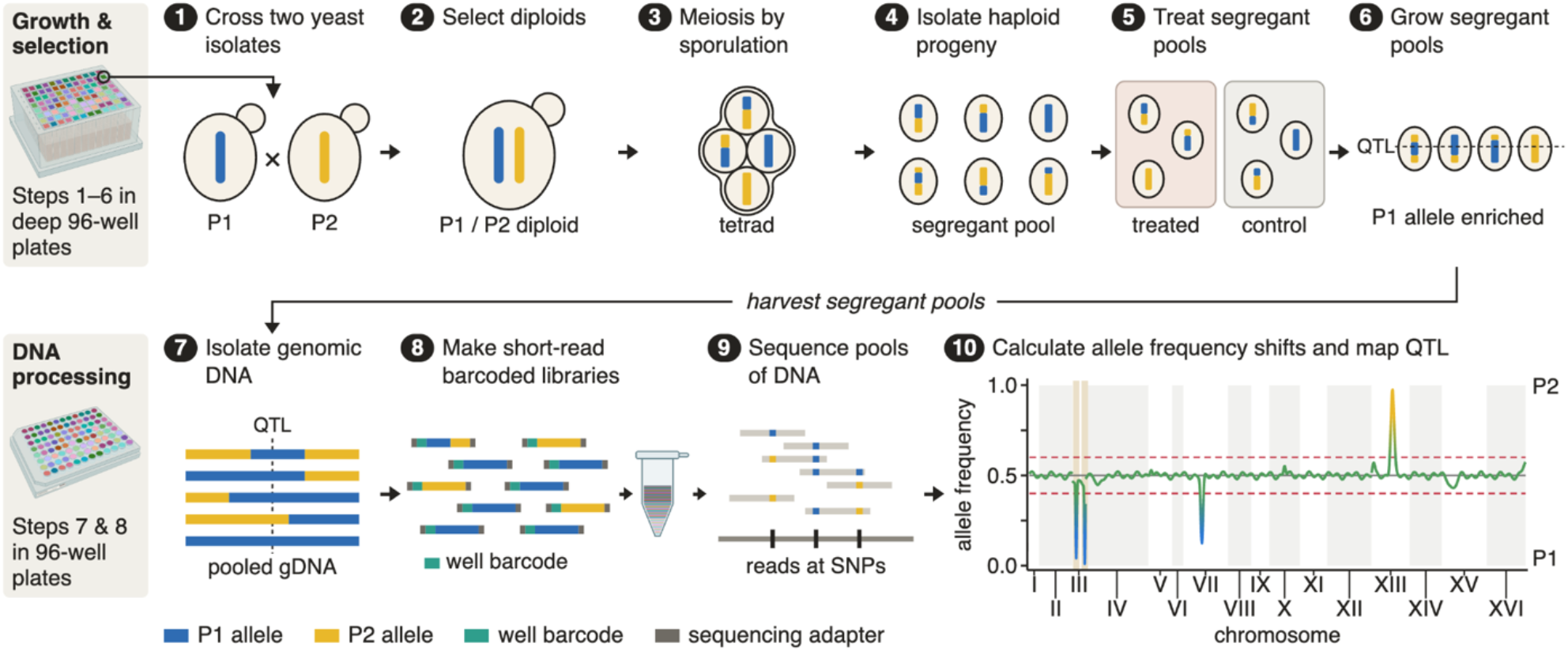
Liquid X-QTL method overview. Top: Steps 1-6 are carried out in 96-deep-well plates. 1) Two genetically distinct yeast isolates are crossed. 2) Diploids are selected with constitutively expressed drug markers integrated at *HO* in both parents. 3) Diploids are starved to induce sporulation, creating four meiotically divided haploid spores in a tetrad. 4) Tetrad asci are digested and haploid progeny are recovered, then selected for a single mating-type using a mating-type specific promoter expressing a drug marker, creating pools of segregant progeny. 5) Segregant pools are grown in control media or matched treated media. 6) Segregant pools are grown to allow sufficient time for some parental alleles to become enriched. Bottom: Steps 7 & 8 are carried out in 96-well PCR plates. 7) Segregant pools are frozen, and genomic DNA is extracted. An example of what 5 individuals may look like at a given allele is displayed below the step heading, where each blue and yellow track represents an individual’s genome in one region. In this example, there is some enrichment for the P1 allele. 8) Genomic DNA is sheared, barcoded, and amplified in each well. All wells are then combined into a single DNA library pool. 9) Our DNA library is sequenced. Reads are aligned to the reference genome, and SNPs are counted across the whole genome for parental alleles P1, or P2 at informative sites. 10) Allele frequencies can then be visualized per well, showing QTL associated with growth, in the case of a control. Our experiment has internal controls at Chromosome 3 and 4 that indicate if a well has been selected properly at step 4. All other regions of the genome where allele frequencies shift may be related to growth in the media, or a growth advantage in a treatment condition conferred by one of the two parents.

First, we replaced the auxotrophic (Tong *et al*. 2001; Ehrenreich *et al*. 2010) and fluorescent (Treusch *et al*. 2015) mating-type selection markers with drug-resistance markers, which work in prototrophic isolates and do not require a cell sorter. We replaced *HO* with the integrating cassette GG55 (*HO* homology arm *Ste2:HygR Ste3:NatR PhleoMX HO* homology arm), which consists of hygromycin resistance driven by the *MAT*a-specific *STE2* promoter, nourseothricin resistance driven by the *MAT*α-specific *STE3* promoter, and a constitutively expressed phleomycin-resistance marker.

Second, we integrated GG55 into a single common parent, the yeast isolate BBQ *MATα HOΔ*::*GG55 Ste2:HygR Ste3:NatR PhleoMX* (hereafter BBQ), which sporulates at close to 100% efficiency (Khaiwal *et al*. 2025), so that each cross yields as many segregants as a fixed culture volume allows. Most of the parental isolates we crossed to BBQ are included in the 1,011 yeast genomes collection (Peter *et al*. 2018) and carry *KanMX* in place of *HO*.

Third, we grew every culture in high-cell-density (HCD) medium (Roberts *et al*. 2020), which supports dense growth in 1 mL, so that selection and growth can take place in deep-well plates. We then isolated DNA from each well, used it to prepare a barcoded short-read library, and then combined the libraries from all the wells for one sequencing run (Materials and Methods).

The workflow in each well works as follows. First, we cross BBQ to a target strain that carries *HOΔ::KanMX*. We then select for diploids in G418 and phleomycin. We sporulate the resulting culture, digest the tetrads, and briefly recover the digested culture, which mostly consists of haploid progeny. We next select haploid segregants of one mating type by using the marker driven by the chosen mating-type specific promoter. In the experiments described here, we selected MATα segregants by nourseothricin selection with the MATα-specific promoter *Ste3:NatR*. All steps take place in liquid culture. The final pool of haploid progeny is then grown in a control condition or selective condition of interest, followed by DNA isolation, shearing, and barcoding. Libraries from 384 wells are pooled for a single sequencing run (Figure 1).

### Thirty-six crosses in four conditions on four plates

We crossed 40 *MAT*a isolates to BBQ in one 96-well plate (Table S9), with each cross represented in at least one well. Two of the 40 isolates turned out to be duplicates of another isolate, and two were absent from the 1,011 sequenced yeast isolate collection and so did not have parental genotypes with which to call segregating sites, leaving 36 crosses for subsequent analyses (Materials and Methods). We took every cross through mating, diploid selection, sporulation and random-spore preparation in deep-well plates and selected *MAT*α segregants in nourseothricin. We then grew each segregant pool in HCD medium (glucose), HCD with 2% acetate replacing glucose, HCD with 1 M NaCl, or HCD with 20 mM caffeine. One 96-well plate was used for each growth condition, and pools were grown for 48 hours, backdiluted and grown for a further 48 hours in the same medium. Acetate was selected as a nonfermentable carbon source (Turcotte *et al*. 2010). We extracted DNA from the 96-hour timepoint for library construction except for the acetate cultures, which did not grow after the second passage, and we extracted DNA from the 48-hour cultures instead. Sodium chloride and caffeine responses were mapped previously by X-QTL (Treusch *et al*. 2015), acetate response was not, and glucose cultures enabled us to control for general genetic effects on growth responses in HCD. We sequenced DNA from all 384 individually barcoded wells as a single library in one run at a target depth of 30×.

Of the 384 wells, 181 yielded usable sequencing data for downstream analyses. 119 wells yielded no sequencing data, and 84 failed one of several quality filters (Materials and Methods; Table 1). The 181 wells with usable data cover 90 of the 144 combinations of 36 crosses and 4 conditions (35 crosses in glucose, 25 in acetate, 13 in caffeine and 17 in sodium chloride), with a median of 2 wells per cross and condition (range 1 to 6; Table 1).

### Fixation of the BBQ allele at *MAT* and *HO*

If haploid selection works as designed, the BBQ allele is expected to fix at two loci in every pool. Because the GG55 cassette at the *HO* locus carries the mating-type selection marker, and the segregant pool is grown in nourseothricin, the only surviving cells should be MATα because the mating-type specific promoter STE3 drives the expression of *NatR*, which confers resistance to nourseothricin. As a result, the BBQ allele of the *HO* locus on chromosome IV, which carries the resistance cassette, is expected to fix. Similarly, because BBQ is the MATα parent in these crosses, the BBQ allele of the MAT locus on chromosome III is expected to fix as a consequence of selection for MATα segregants.

In the absence of selection, random segregation should lead each parental allele to have ∼0.5 frequency across the genome; at a fixed locus, the frequency of the selected parental allele should instead approach 1. Across the 181 segregant pools, the median BBQ allele frequency was observed to be 0.99 at MAT (range 0.91 to 1) and 0.99 at *HO* (range 0.71 to 1) (Figure 2; Table S4), similar to previous X-QTL studies (Ehrenreich et al. 2010; Albert et al. 2014). The MAT locus was fixed for the BBQ allele in all 181 segregant pools. The *HO* locus was fixed in 178; the three exceptions (BBQ allele frequency 0.71 to 0.83) occurred in 3 replicate wells seeded from a single segregant preparation of cross AAM×BBQ. These results confirm that selection of MATα haploid segregants worked well in the 96-well format. The *MAT* and *HO* loci were masked in subsequent QTL mapping analyses (Materials and Methods).

**FIGURE 2.**
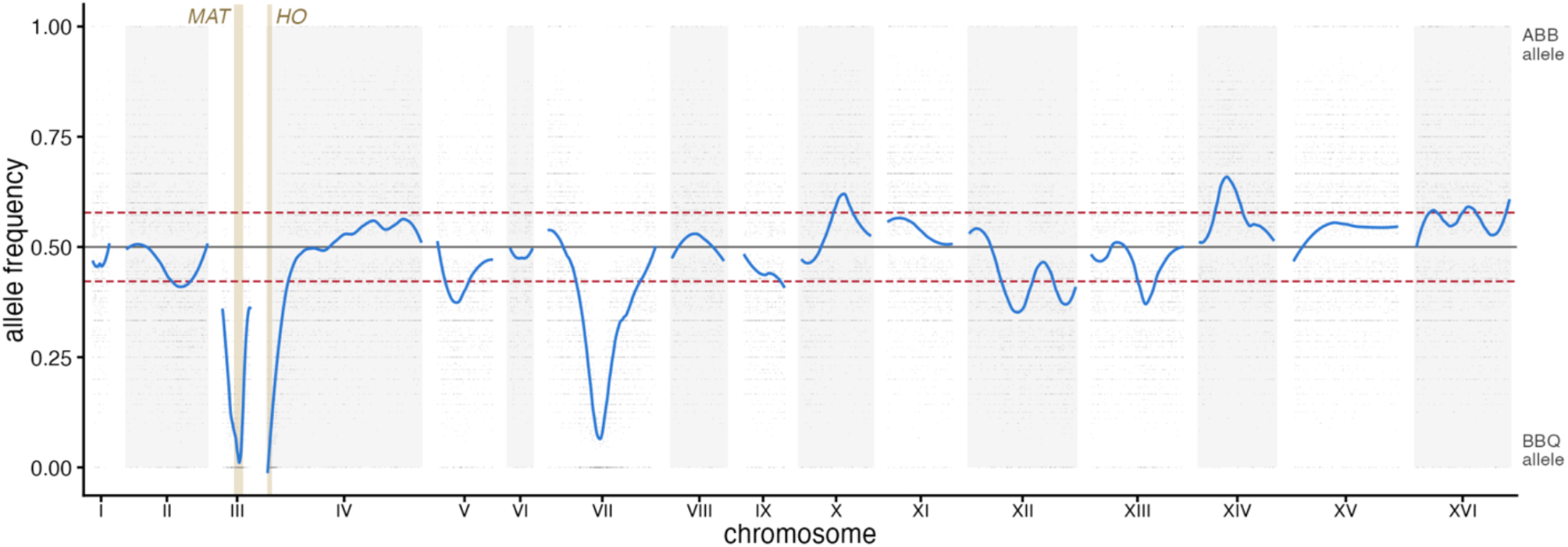
Segregant pools return fitness and condition specific QTL. (A) Allele frequency across the genome from ABBxBBQ’s segregant pool in glucose (well B02). Grey points are per-site allele frequency estimates, and the blue line is the smoothed fit. An allele frequency of 1 indicates the ABB allele fixed and 0 that the BBQ allele fixed. The dashed red lines mark the smallest effect that reaches LOD 5 in this pool. Tan bands are the mating-type region and the *HO* locus, expected to fix to BBQ alleles in each cross.

### QTL Mapping

We carried out QTL mapping for each well. At single nucleotide sites that differed between the two parents of a cross (informative sites; Materials and Methods), we counted the reads carrying each parental allele and smoothed the allele frequency along each chromosome. At loci that affect fitness in the tested growth condition, the frequencies of the two alleles will deviate from 0.5 in the direction of the parent that carries a variant that increases growth (Figure 2). We converted the allele frequency skew at each position into a LOD score (log10 of the odds ratio) and used a LOD threshold of 5 to call significant QTL (Materials and Methods). We observed 1,472 QTL in 179 of the 181 segregant pools, representing 35 of the 36 crosses (Table 2), with a median of 6 QTL per pool (range 1 to 53). The BBQ allele was favored at 74.3% of QTL (70.9% excluding the chromosome VII QTL described below).

QTL recur across crosses. When grouped within each condition, QTL recur in 74 overlapping regions in glucose, 77 in acetate, 88 in caffeine and 89 in sodium chloride (Table 3). Only one region recurs in nearly every cross. A region on chromosome VII (Table S4) skews toward the BBQ allele in 34 of 35 crosses and carries a QTL in 170 of the 179 segregant pools with QTL (95.0%), in all four conditions. The BBQ allele is favored at all 170 of these QTL, at a median skew of 0.390. Future work will identify the variant underlying this BBQ allele.

QTL detected in segregant pools grown in HCD with glucose correspond to general genetic effects on growth, rather than to loci that influence growth in response to a specific stressful condition. To isolate the QTL that specifically influence growth in sodium chloride, caffeine, or acetate we computed the difference in allele frequency between each pool grown in the condition of interest and the same pool grown in the glucose control condition (Figure 4; Table 2), as in previous X-QTL experiments (Ehrenreich *et al*. 2010; Torabi and Kruglyak 2012; Treusch *et al*. 2015). This difference was converted to a LOD score as above, and used to call 314 condition-specific QTL in 75 segregant pools of 24 crosses.

Condition-specific QTL recur across crosses less often (Figure 3). When grouped within each condition, condition-specific QTL recur in 43 overlapping regions in acetate, 79 in caffeine and 56 in sodium chloride (Table 8). Each region recurs in a mean of 1.4 to 1.6 crosses, against 2.3 to 4.4 crosses per region before the contrast (Tables 3 and 8). The regions are also narrower, with median widths of 20.0 to 41.4 kb (Table 2).

**FIGURE 3.**
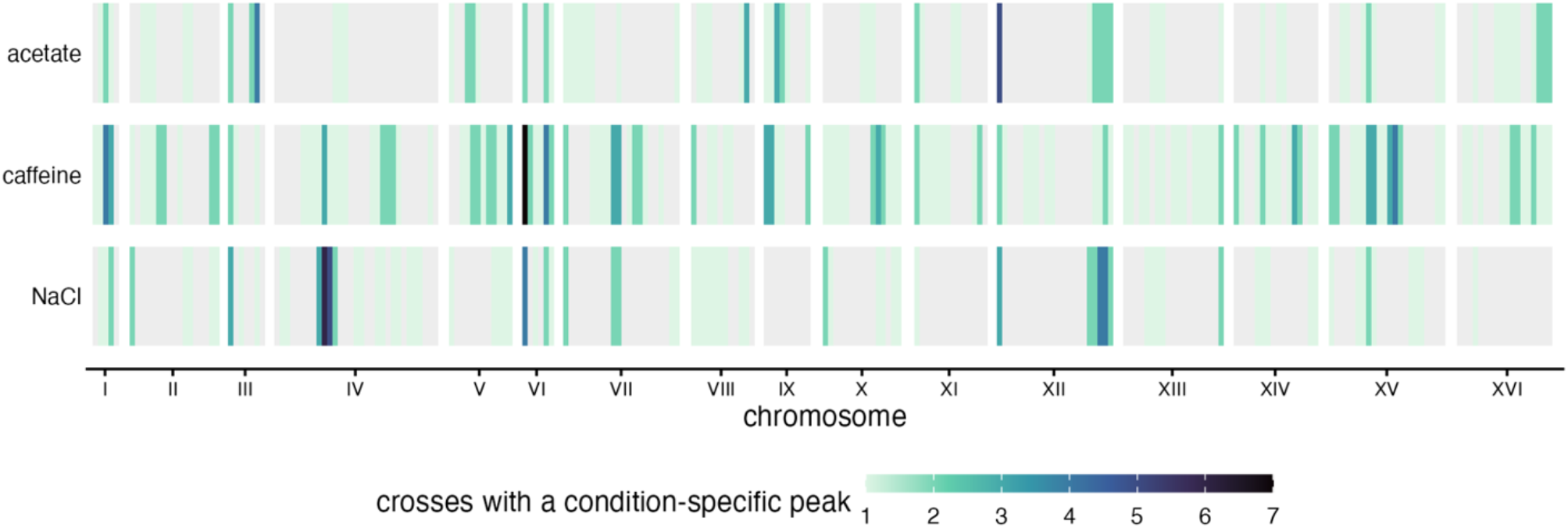
Condition-specific QTL per genomic bin across all crosses. The genome is divided into 50 kb bins and each bin is colored by the number of distinct crosses with a condition-specific QTL called at LOD 5 or above whose interval falls within that bin. Each condition is labelled in the y-axis. Bins covered by no QTL are light grey. QTL inside the mating-type region on chromosome III and the *HO* locus on the left arm of chromosome IV are removed before binning. A QTL peak called just outside one of those regions can still contribute its interval to neighboring bins.

### Comparison with previously mapped QTL for sodium chloride and caffeine resistance

Treusch *et al*. (2015) used X-QTL to map 37 loci for sodium chloride tolerance and 23 for caffeine tolerance in round-robin crosses between twelve isolates (Table S3). Although these isolates are distinct from those used in the present study, some QTL may be detected in both studies as a consequence of causal variants that are common in the yeast population or of recurrent mutations in the same causal genes. Sodium chloride resistance in Treusch *et al*. was dominated by the *ENA* locus on chromosome IV, which was detected in 11 of the 12 crosses (Table S1). The *ENA* genes encode sodium efflux pumps (Haro *et al*. 1991; Márquez and Serrano 1996). European isolates carry an introgressed *ENA* allele from *S. paradoxus* in variable copy number, while other isolates carry a single divergent *ENA6* gene (Daran-Lapujade et al. 2009; Warringer et al. 2011). Both the introgressed allele and its copy number are associated with salt tolerance (Warringer et al. 2011; Wilkening et al. 2014; Treusch et al. 2015). While 36 crosses went into the experiment, sodium chloride and caffeine had large losses of wells during sample preparation, yielding 17 and 13 usable crosses, respectively. We detected a sodium chloride QTL overlapping the *ENA* locus in 5 of 17 usable condition specific crosses (Table 1; Table 9). The BBQ allele was favored in four crosses (Figure 4A) and the alternate-parent allele in one cross (Figure 4B), consistent with known allelic heterogeneity (Daran-Lapujade *et al*. 2009, Warringer *et al*. 2011,Wilkening *et al*. 2014, Márquez and Serrano 1996, Treusch *et al*. 2015). Treusch *et al*. (2015) identified *TOR1* and *WHI2* as causal genes underlying caffeine QTL. We detected a caffeine QTL overlapping *WHI2* in 3 of 13 crosses and a QTL overlapping *TOR1* in one cross (Table 9). Across the two conditions, 24 of the 60 published QTL overlapped a QTL in at least one of our crosses (12 of 23 in caffeine and 12 of 37 in sodium chloride; Table 10). Our study also identified many new condition-specific QTL. 64 of the 79 caffeine QTL regions do not overlap QTL reported by Treusch *et al*. (2015), and neither do 36 of the 56 sodium chloride QTL (Table 8).

**FIGURE 4.**
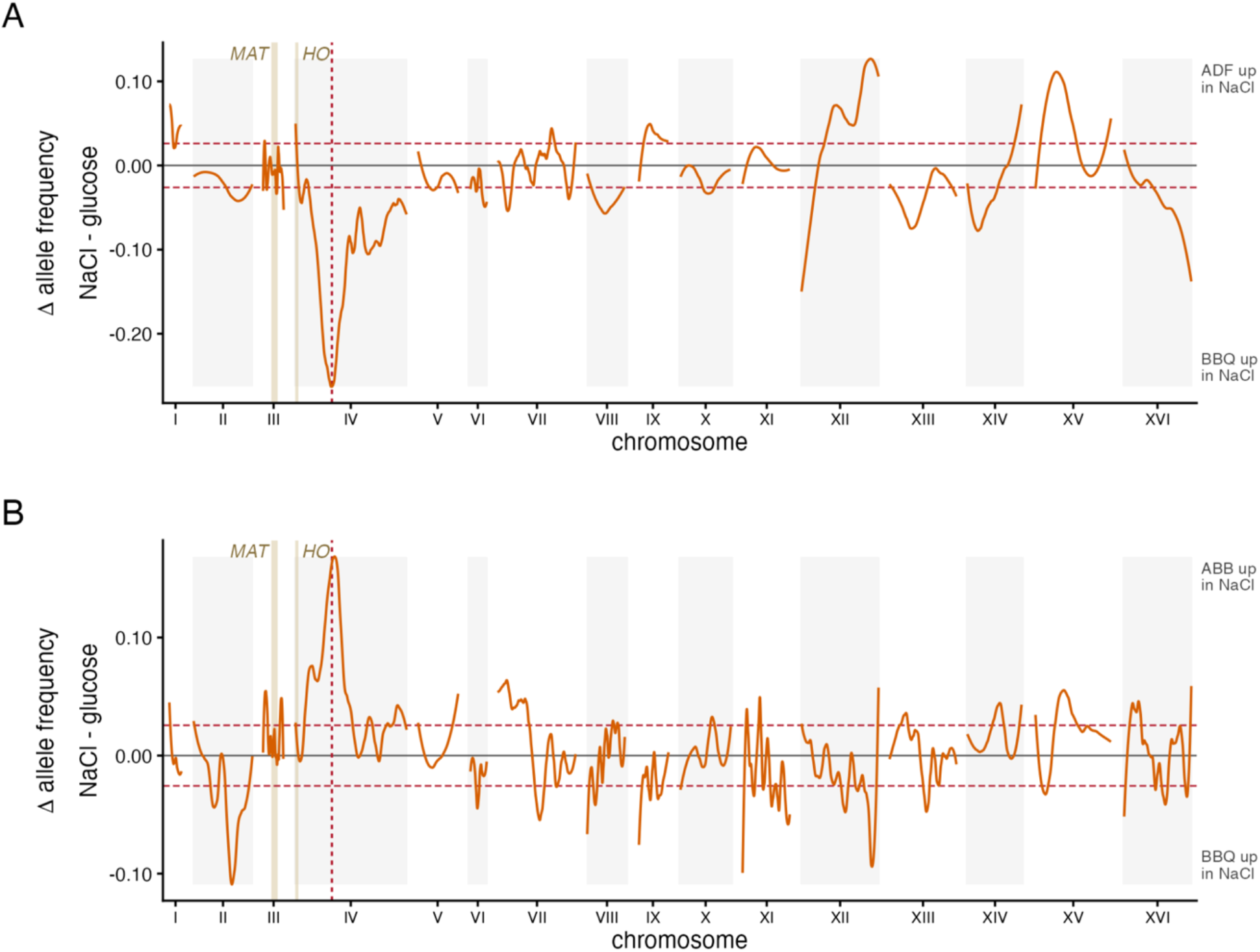
Detection of published loci. (A) The effect of sodium chloride on the difference in allele frequency across the genome from ADFxBBQ’s segregant pools. The y axis is the sodium chloride allele frequency minus the glucose allele frequency, so positive values mark positions where the ADF allele was enriched under sodium chloride and negative values mark positions where the BBQ allele was enriched. The dashed red lines mark the smallest difference that reaches LOD 5. The ENA locus on chromosome IV at 509,600 is marked. (B) The effect of sodium chloride on the difference in allele frequency across the genome from ABBxBBQ’s segregant pools. The y axis is the sodium chloride allele frequency minus the glucose allele frequency, so positive values mark positions where the ABB allele was enriched under sodium chloride and negative values mark positions where the BBQ allele was enriched. The dashed red lines mark the smallest difference that reaches LOD 5. The ENA locus on chromosome IV at 509,600 is marked.

## DISCUSSION

We adapted X-QTL to 96-well plates and mapped QTL in 35 crosses in four conditions from 384 wells in one sequencing run. We detected loci previously mapped by conventional X-QTL, including the *ENA* locus, which influences growth in sodium chloride, and the *WHI2* and *TOR1* loci, which influence growth in caffeine. We obtained usable data from 181 of the 384 wells; the rest either did not yield DNA or did not pass sequencing data quality filters (Table 1). Future improvements in library yield will increase the throughput and robustness of this method.

Two features of the study design influenced the results of QTL mapping: the use of a single common parent for all crosses, and few replicate wells per cross and condition. Because every cross shares the parent strain BBQ, BBQ alleles that increase relative fitness segregate in multiple crosses, whereas alleles unique to one of the other parent strains segregate only in the cross between BBQ and that strain. We observed that BBQ alleles were favored at 74.3% of detected QTL, and one chromosome QTL at which the BBQ allele was favored was detected in 34 of 35 crosses. The use of a common parent also limits the comparison with prior studies, such as that of Treusch *et al*. (2015): we can only detect loci at which the BBQ strain differs from one of the other parent strains.

Integrating the GG55 cassette into additional high-sporulation strains will allow a greater range of genetic variation to be accessed. Each cross was also represented by a median of only two replicate wells per condition (range 1 to 6). More wells per cross and condition would increase statistical power at the expense of throughput.

In our new method, each microplate well is a self-contained experiment for a segregant pool in a specific condition. Libraries from at least 384 wells can be sequenced at once, allowing for efficient QTL mapping across a much larger space of crosses and growth conditions. By removing the need for a separate experiment for each cross and condition, our method will power studies of interactions between genetic effects and environments in many genetic backgrounds.

## MATERIALS AND METHODS

### Yeast strains and growth

The common parent BBQ and alternate parents from the 1011 Yeast Genomes Project (Peter et al., 2018) served as the parental backgrounds for all crosses. A complete list of strains used is provided in Table S9. Unless otherwise noted, strains were grown in YPD (1% yeast extract, 2% peptone, 2% glucose) at 30 °C with shaking at 220 rpm in 15 ml culture tubes, or 1000 rpm in 2 ml deep well plates. High Cell Density Media (HCD) was used for some experiments as noted, see the key resource table or Roberts *et al*. (2020) for formulation. Selection plates and liquid cultures contained YPD or HCD supplemented with the appropriate antibiotics: G418 200 µg/mL, nourseothricin (clonNAT) 100 µg/mL, hygromycin B 300 µg/mL, and/or phleomycin 20 µg/mL. Long-term stocks were maintained at –80 °C in 16-25% (v/v) glycerol.

### Magic-marker plasmid and integration at the *HO* locus

The plasmids GG55_Ste2:HygR_Ste3:NatR_PhleoMX (7,497 bp; [ADDGENE # — TO BE FILLED IN]) and GG56_Ste2:KanR_Ste3:NatR_PhleoMX (6740 bp; [ADDGENE # — TO BE FILLED IN]) were constructed by Golden Gate assembly and carry two mating-type-resolvable selectable markers flanked by *HO* homology arms: *STE2pr-KanR/HygR* (G418, and hygromycin resistance markers, respectively, under a *MATa*-specific promoter), *STE3pr-NatMX* (nourseothricin resistance under a *MATα*-specific promoter), and a constitutively expressed *PhleoMX* marker (phleomycin resistance) for selection of diploids and strains integrated with GG55 or GG56 regardless of mating type. GG55 and GG56 have 300 bp *HO* homology arms that are PCR amplified to create a linear integration cassette for transformation. The cassette was integrated at the endogenous *HO* locus by lithium acetate transformation as described below; integration disrupts *HO* and thereby prevents mating-type switching, while the *STE2* and *STE3* promoters subsequently enable mating-type-specific selection of haploid progeny after sporulation.

### Yeast transformation

Parental strains were transformed with integration cassette DNA (linear GG55 or GG56 amplified with rGNB047 and rGNB048) using a standard lithium acetate, PEG, single-stranded carrier DNA protocol. Overnight cultures grown in 2 mL YPD at 30 °C with shaking were back-diluted into 50 mL YPD to a starting OD_600_ of 0.1 and grown to mid-log phase (OD_600_ 0.2–0.8, ∼3–4 hours at 30 °C, 220 rpm). Cells were harvested by centrifugation (3,000 × g, 3 min), washed in 10 mL 100 mM LiOAc, pelleted, and resuspended in 0.5–1 mL LiOAc. For each transformation, 100 µL of cell suspension was combined with 1-10 µL linear DNA amplified by PCR (≤100 ng), 10 µL boiled salmon sperm carrier DNA, and 280 µL 50% (w/v) PEG 3350, vortexed, and incubated on the benchtop for 30 min. 43 µL DMSO was added, and reactions were heat-shocked at 42 °C for 10 min. Cells were pelleted at 1,500 × g for 2 min, washed once in 0.5 mL sterile water, and plated directly onto YPDA. Yeast were recovered for 18 hours at 30 °C, then replica plated on selective medium YPDA + Phleomycin.

### Parental BBQ haploid creation and sporulation

To build the parental haploid line with the magic marker cassette we did the following. Parental BBQ *HO/HOΔ*::*GG55 Ste2:HygR Ste3:NatR PhleoMX* strains were sporulated using a standard sporulation protocol. Briefly, the strain indicated above was grown overnight in a 15 ml culture tube with 2 ml YPD + Phleomycin. The overnight culture was collected, and 0.5 mL was washed in water and resuspended in 5 ml SPO media in a 15 ml culture tube. Tubes were placed on a rotary wheel, and cell suspensions were slowly rotated for 2-7 days. Tetrad formation was checked daily. Upon detections of tetrads, sporulated cultures were pelleted at 500 xg for 3 min, washed in 0.5 mL water, and resuspended in 200 µL water per well. Zymolyase 20T (40 µL of a 500 U/mL stock; 20 U per well) was added, and the digestion was incubated at room temperature for 20–40 min with periodic monitoring. The reaction was quenched by addition of 300 µL 1 M sorbitol, and cells were pelleted (3,500 rpm, 3 min), the supernatant removed, and the pellets washed in 0.5 mL water.

After a second wash, cells were spread on a YPD plate and tetrads were dissected on a Singer tetrad dissection microscope. Tetrads were then genotyped and phenotyped for mating-type and integrating cassette retention.

### Verification of cassette integration

Single transformant colonies were screened for integration at *HO* by colony PCR. Eight colonies were tested per transformation. PCR mixtures (50 µL) contained 25 µL OneTaq 2X Master Mix (New England Biolabs), 1 µL each of forward primers rGNB146 (endogenous *HO* exon 1) and rGNB174 (MX integration), 1 µL reverse primer rGNB145 (downstream of the *HO* right arm), 1 µL 25% Triton X-100, and nuclease-free water to 50 µL. A small piece of yeast colony (∼1–2 pipette-tip diameters) was added directly to each reaction using a plastic p10 tip and swirled to dislodge; the same tip was used to streak a numbered sector of a YPDA + phleomycin plate, which was incubated at 30 °C for 1–2 days to recover the corresponding clone. Thermocycling: 94 °C for 30 s; 30 cycles of 94 °C for 15 s, 52 °C for 30 s, 68 °C for 1 min; final extension at 68 °C for 5 min.

Products were resolved on 1% agarose gels at 100 V. Integration of either GG55 or GG56 cassette yielded a 495 bp amplicon (rGNB145 + rGNB174), whereas the endogenous *HO* locus yielded a 1,015 bp amplicon (rGNB145 + rGNB146); the triplex reaction generated both products in heterozygotes and at single-locus resolution distinguished integrants from non-integrants. Colonies with confirmed integration were used to prepare glycerol stocks.

### Verification of haploid mating type

Mating-type-selected haploids and clones were genotyped by multiplex PCR using primers MAT-a (5′-CAATGATTAAAATAGCATAGTCGG-3′), MAT-α (5′-CAGCACGGAATATGGGAC-3′), and MAT-R (5′-GGTGCATTTGTCATCCGTC-3′) (Bradbury et al., 2006). Reactions (25 µL) contained 2 µL of overnight YPD culture (diluted 1:50) as template and were performed with OneTaq. Cycling: 94 °C for 10 min; 35 cycles of 94 °C for 25 s, 55 °C for 25 s, 72 °C for 90 s. Products (*MATa*, 489 bp; *MATα*, 466 bp) were resolved on a 3% agarose gel run at 100 V for 4 hours in sodium borate buffer (10 mM NaOH, 36 mM boric acid) stained with ethidium bromide. The *HO* locus was additionally sequenced in a subset of confirmed haploids to verify cassette structure via sanger sequencing.

### 96-well mating

Diploids were generated using a 96-well mating scheme in which a single common parent was crossed against an arrayed panel of partners. The common parent was grown overnight in 3 mL YPD at 30 °C, 220 rpm. The partner panel was inoculated from a 96-well glycerol stock plate into a fresh 96-well plate containing 100 µL YPD per well using a sterile pin replicator and grown overnight at 30 °C, 220 rpm. For mating, 20 µL of each partner culture and 20 µL of the common-parent culture were combined into a new 96-well plate containing 80 µL YPD per well and incubated at 30 °C overnight without shaking; a visible pellet was observed at 24 hours. Mating mixtures were resuspended by vortexing at 1,000 rpm for 30 s and pinned into a deep-well plate containing 1 mL YPD + G418 + phleomycin per well to select for diploids carrying both parental cassettes. After overnight growth at 30 °C, 1,000 rpm, cultures were pinned a second time into fresh double-selection medium to further dilute residual haploid parents. Glycerol stocks of mated diploids were prepared by mixing 80 µL culture with 80 µL 50% glycerol.

### 96-well sporulation

Diploid cultures from the mated plate were transferred (750 µL per well) to a new deep-well 96-well plate, pelleted at 3,700 rpm for 1 min, washed once in 0.5 mL YPA pre-sporulation medium (1% yeast extract, 2% peptone, 2% potassium acetate), and resuspended in 750 µL YPA. For sporulation, 300 µL of the diploid suspension was combined with 700 µL YPA in a deep-well 96-well plate and incubated overnight at 30 °C, 800 rpm. The following morning, cells were pelleted (3,700 rpm, 1 min), the YPA supernatant was removed, and pellets were resuspended in 1 mL SPO medium (1% potassium acetate) and returned to the same incubator and agitation conditions. Sporulation efficiency was monitored by hemocytometer counts of tetrads and asci relative to vegetative cells starting at 48 h; cultures were harvested when sporulation reached 12–90% (strain-dependent).

### 96-well random spore preparation

Sporulated cultures were pelleted at 3,500 rpm for 3 min, washed in 0.5 mL water, and resuspended in 200 µL water per well. Zymolyase 20T (40 µL of a 500 U/mL stock; 20 U per well) was added, plates were mixed at 1,000 rpm for 15 s, and the digestion was incubated at room temperature for 20–40 min with periodic monitoring. The reaction was quenched by addition of 300 µL 1 M sorbitol, and cells were pelleted (3,500 rpm, 3 min), the supernatant removed, and the pellets washed in 0.5 mL water. After a second centrifugation, pellets were resuspended in 0.5 mL YPD and allowed to recover at 30 °C, 1,000 rpm for 4 h. Cultures were either archived as glycerol stocks (500 µL culture + 500 µL 50% glycerol, –80 °C) or split (250 µL per well) into two daughter plates containing YPD + G418 or Hyg (selecting *MATa* segregants via *STE2pr-KanR or HygR*) or YPD + nourseothricin (selecting *MATα* segregants via *STE3pr-NatR*).

### Treatment of haploid segregant pools

Haploid segregant pools were thawed fully from glycerol stocks, and 250 µl was removed (∼1/6 of full culture stock), washed in HCD, resuspended in 1 ml HCD, then grown overnight at 30°C 1000 RPM. Each treatment condition (Caffeine 20 mM, NaCl 1 M, 2% Acetate instead of glucose, HCD control) was seeded at 16 µl into 984 µl and grown for 48 h. From those segregant pools 12 µl was seeded into the exact same condition in 1 mL for an additional 48 hours to create a bottleneck, with the exception of Acetate, as the 96 hour cultures did not survive. Following treatment, 200 µL aliquots from each plate were transferred to PCR tubes and pelleted; the remainder of each plate was pelleted and frozen at –80 °C for archival storage.

### Genomic DNA isolation by 96-well ethanol precipitation

Genomic DNA was isolated from harvested cell pellets in 96-well format. Cells from 200 µL of overnight culture (or equivalent treatment endpoint volume) were transferred to a semi-skirted twin.tec PCR plate (Eppendorf, E951020303) seated in a holder plate (Thermo Scientific AB0800), pelleted at 4,000 rpm for 2 min, and the supernatant removed. Pellets were resuspended in 30 µL of Buffer 1 (1 M sorbitol, 0.1 M EDTA, pH 7.5) containing Zymolyase 20T at a working concentration of 13 mg/mL (5.2 µL of 250 mg/mL stock per 100 µL Buffer 1) using the plate-mixer setting on a Eppendorf MixMate shaker. Plates were incubated at 37 °C for 30 min with mixing every 10 min to prevent settling. 30 µL of Buffer 2 (50 mM Tris-HCl, 20 mM EDTA, 1% w/v SDS, pH 7.4) was added, plates were sealed with foil and incubated at 65 °C for 5 min, then briefly centrifuged (4,000 rpm, 30 s). Proteins and SDS were precipitated by addition of 24 µL 5 M potassium acetate per well, mixing, and incubation at –20 °C for 5 min (without freezing the pellet). Plates were centrifuged at 4 °C, 4,000 rpm for 30 min. Cleared supernatants (∼50 µL) were transferred using an INTEGRA VIAFLO into a fresh plate containing 75 µL isopropanol per well, mixed by pipette, and incubated at room temperature for 15 min. DNA was pelleted at 15 °C, 4,000 rpm for 20-40 min; supernatants were removed and pellets washed with 150 µL ice-cold 70% ethanol (without mixing) and centrifuged at 4 °C, 4,000 rpm for 30 min. Ethanol was removed and pellets air-dried at 60 °C for 60–90 min before resuspension in 100 µL nuclease-free water at 55 °C for 15 min. Yields were quantified by Qubit dsDNA HS Assay (typical range 5–80 ng/µL). DNA was diluted to 1 ng/µl for downstream processing.

### Tn5 tagmentation and library preparation

Sequencing libraries were prepared using a homebrew Tn5 tagmentation protocol, with transposase pre-charging and tagmentation conditions adapted from Picelli *et al*. (2014) and Stern (2017) and further modified by the Kruglyak lab. All steps were performed in 96-well format. Briefly:

#### Adapter annealing

Tn5 mosaic-end adapters were annealed in two separate reactions in PCR tubes: Reaction 1 contained 10 µL Tn5ME-A (100 µM) + 10 µL Tn5MErev (100 µM) + 80 µL Reassociation Buffer (10 mM Tris-HCl pH 8.0, 50 mM NaCl, 1 mM EDTA); Reaction 2 contained 10 µL Tn5ME-B (100 µM) + 10 µL Tn5MErev (100 µM) + 80 µL Reassociation Buffer. Annealing was performed in a thermal cycler programmed for: 95 °C for 10 min; 90 °C for 1 min; 60 cycles of stepwise –1 °C/cycle; 4 °C hold. Annealed adapters (10 µM final each) were stored at –20 °C; working dilutions (1 µM) were prepared in nuclease-free water. (Picelli et al. 2014)

#### Tn5 transposase preparation and pre-charging

Purified Tn5 transposase (1.7 mg/mL stock in 25 mM Tris-HCl pH 7.5, 800 mM NaCl, 0.1 mM EDTA, 1 mM DTT, 50% glycerol) was diluted to a working stock (672 µg/mL) by combining 20 µL of the 1.7 mg/mL stock with 24 µL 100% glycerol and 6.6 µL freshly prepared TN5 Dialysis Buffer (100 mM HEPES pH 7.2, 0.2 M NaCl, 0.2 mM EDTA, 2 mM DTT, 0.2% Triton X-100, 20% glycerol). The working stock was further diluted 1:10 in Reassociation Buffer to yield the Working Dilution (67.2 µg/mL). For pre-charging, equal volumes (10 µL each) of the Working Dilution, 100% glycerol, and 1 µM annealed adapter pair were combined in a final volume of 41 µL (34.4 µg/mL Tn5 with 243 nM adapters), and the mixture was incubated at 37 °C for 30 min.

#### Tagmentation

Pre-charged Tn5 was further diluted 1:10 in the tagmentation master mix to yield a final concentration of ∼0.34 µg/mL Tn5 with 24.3 nM adapters in the reaction; this dilution was empirically determined to avoid over-tagmentation of xQTL pool libraries (the standard concentration of ∼2 ng/µL described in earlier implementations consistently over-tagmented our libraries). The tagmentation reaction (10 µL final per sample) contained 0.1 µL diluted pre-charged Tn5, 2 µL 5× TAPS-DMF buffer (50 mM TAPS pH 8.5, 25 mM MgCl_2_, 50% DMF), 2 µL 40% PEG 8000, 4.9 µL water, and 1 µL genomic DNA at 1 ng/µL. 9 µL of master mix was dispensed per well using an electronic multichannel pipette (multi-dispense mode) into a labeled 96-well PCR plate; 1 µL of genomic DNA was added and mixed by repeated pipetting. Plates were transferred to a pre-warmed thermal cycler and incubated at 55 °C for 3 min. Reactions were stopped immediately by addition of 2.5 µL 0.2% SDS per well, mixed, and incubated at 55 °C for an additional 7 min.

#### Indexing PCR

Tagmented DNA was indexed by PCR using dual-indexed Illumina i5/i7 primers (IDT for Illumina Nextera DNA UD Indexes Set A-D). Each 20 µL reaction contained 10 µL KAPA HiFi HotStart 2X ReadyMix, 1 µL DMSO, 1 µL water, 2 µL i5/i7 primer mix, and 6 µL tagmented DNA. Reactions were assembled by dispensing 12 µL of master mix per well, followed by 6 µL of tagmented DNA and 2 µL of indexed primer mix. Cycling: 72 °C for 3 min; 95 °C for 30 s; 20 cycles of 98 °C for 20 s, 58 °C for 15 s, 72 °C for 30 s; final extension at 72 °C for 3 min; 10 °C hold.

#### Pooling, cleanup, and size selection

19 µL from each indexed well was pooled in a nuclease-free reservoir, mixed gently to avoid bubble formation from residual detergents, and transferred to a microcentrifuge tube. The pooled library was subjected to a two-step KAPA Pure Bead cleanup and size selection following manufacturer guidelines (Roche/KAPA Biosystems, KK8001). First, a 0.8× single-sided cleanup was performed on 100 µL of pooled library by addition of 80 µL KAPA Pure Beads, incubation at room temperature for 5–15 min, two 80% ethanol washes, and elution into 100 µL 10 mM Tris-HCl pH 8.0; this step is essential to remove PEG carried over from the tagmentation buffer prior to size selection. Second, a 0.5–0.7× double-sided size selection was performed by adding 50 µL KAPA Pure Beads to the 100 µL cleaned library (first cut, excluding fragments >1 kb), transferring the supernatant, and adding an additional 20 µL beads (second cut, retaining fragments 300–750 bp), followed by 2 × 200 µl 80% ethanol washes and elution in 50 µL 10 mM Tris-HCl pH 8.0.

#### Library quantification

Final libraries were quantified by Qubit dsDNA Assay and profiled on an Agilent TapeStation HS D1000 to determine the mean fragment size. Absolute molar concentration of sequenceable molecules was then determined by quantitative PCR using the KAPA Library Quantification Kit (Illumina, Roche/KAPA Biosystems) against the kit-supplied six-point DNA standard series (20 pM to 0.0002 pM). Libraries were serially diluted (1:10 through 1:10^6^) in 10 mM Tris-HCl + 0.1% Tween-20, pH 8.5 in DNA LoBind 96-well plates; the three dilutions falling within the linear range of the standard curve (typically 1:1,000–1:100,000) were assayed in technical duplicate alongside two replicates of each standard and two no-template controls. Reactions (10 µL) contained 6 µL of KAPA SYBR FAST qPCR Master Mix with primer premix and 4 µL of diluted library or standard. qPCR was performed on a BioRad CFX Opus 96 Real-Time PCR System with the following program: 95 °C for 5 min; 35 cycles of 95 °C for 30 s, 60 °C for 45 s (with data acquisition); followed by a 65–95 °C melt curve. Library concentrations were calculated from the standard curve and adjusted for mean fragment size to obtain final molarity.

### Sequencing

Libraries were diluted to 2 nM in nuclease-free water in a minimum volume of 30 µL and sequenced on an Illumina NovaSeq X using a 1.5 B 300 cycle reagent kit with 2 × 150 bp paired-end reads.

### Sequencing data analysis

#### Read processing and parental identification

Reads were aligned to the sacCer3 reference assembly with bwa mem 0.7.19 (Li 2013) and sorted by genomic position with sambamba 1.0.1 (Tarasov *et al*. 2015). Allele counts were quantified at known SNPs in the 1,011 Yeast Genomes panel (Peter et al. 2018) with GATK 4.6.2.0 ASEReadCounter (McKenna *et al*. 2010; Castel *et al*. 2015) in a Nextflow 25.10.4 pipeline (Di Tommaso *et al*. 2017) for reads with a minimum mapping quality of 20, yielding reference and alternate counts per site per pool. For each cross, we retained sites where the two parents were homozygous for different alleles and assigned reads to their respective parental alleles.

Alignment QC used samtools 1.24 (Danecek *et al*. 2021). Allele frequencies, contrasts, peak calling and the downstream tests were computed in R 4.3.3 (R Core Team 2024) with xQTLStats 0.2 (J. S. Bloom, github.com/joshsbloom/xQTLStats, commit 080a156), data.table 1.17.8 (Barrett *et al*. 2025), IRanges 2.36.0 and GenomicRanges 1.54.1 (Lawrence *et al*. 2013). Tables and figures were produced in R 4.5.3 (ggplot2 4.0.2, Wickham 2016; patchwork 1.3.2, Pedersen 2025) and Python 3.10. We ran all other plotting and calculations on the Hoffman2 cluster at UCLA. All code is available at https://github.com/brunig/microscale-xqtl.

Because sequencing data revealed inconsistencies with the original plate map, we reassigned parents directly from sequencing data. BBQ was fixed as one parent, and candidate alternate parents were evaluated by comparing observed allele counts with their expected frequencies under each parental genotype. Data from the four conditions sharing each well position were combined for identification. Candidate assignments were further evaluated using allele-frequency distributions and local residual variation. Of 384 experimental wells 24 with unresolved parentage were excluded (Table S10).

#### Quality control and allele-frequency estimation

Pools were excluded from QTL analysis if they failed sequencing quality control, had unresolved parentage, insufficient coverage, or unreliable allele-frequency estimates. The minimum median sequencing depth was 2×, and pools were additionally excluded if their mean depth was ≤1×, more than 1% of smoothed allele frequencies fell outside the range [0,1], or the median absolute allele-frequency skew exceeded 0.05. Overall, 181 pools passed quality control, 84 were flagged, and 119 had no usable data.

The median-depth threshold was initially set to 5× and subsequently reduced to 2× after evaluating QTL concordance between replicate wells of the same cross and condition.

Allele counts were aggregated into 500-bp blocks, excluding blocks with fewer than 10 reads. Parental allele frequencies were smoothed separately for each chromosome using quadratic LOESS, with the span selected by corrected Akaike information criterion (AICc) (Huang *et al*. 2020).

### QTL identification

We calculated two allele-frequency contrasts: (1) deviation from equal parental representation, defined as AF − 0.5, and (2) the difference between stress-treated and glucose-control pools of the same cross. The latter measures differences between conditions at harvest rather than changes from an initial population, because time-zero samples were not sequenced. For each contrast, we calculated a z statistic using binomial standard errors, with an effective sample-size parameter defined as the smaller of 9,000 and total allele depth divided by 600. Variances were summed for contrasts between two pools. The corresponding two-sided normal probabilities were converted to LOD scores using a one-degree-of-freedom chi-square transformation.

Candidate QTL were called at local maxima exceeding LOD 5. Intervals extended from each peak until the score fell by 10% of its peak value, subject to a minimum width of 20 kb and a maximum of 200 kb. Overlapping intervals within a pool were merged, retaining the higher-scoring peak.

The MAT and HO regions, which were strongly selected during segregant isolation, were excluded from QTL counts and downstream analyses (Table S4).

### QTL grouping and comparison with published loci

Within each condition, overlapping QTL intervals were merged across crosses into loci, following Treusch et al. (2015). For candidate-gene analyses, we instead grouped peak summits within 25 kb and defined consensus regions from the portion of the interval pileup reaching at least half its maximum height, avoiding excessive merging of neighboring loci.

We compared our QTL with the 60 grouped loci reported by Treusch et al. (2015), restricting comparisons to caffeine and sodium chloride. A published locus was considered recovered when its reported peak position fell within one of our QTL intervals for the same condition. Each locus was counted at most once per cross and condition.

We primarily used stress-versus-glucose contrasts to match the control-subtracted design of the earlier study. Enrichment of overlapping loci was evaluated using 1,000 circular-shift permutations of QTL intervals along the concatenated genome, retaining interval number, width, and relative spacing. A recurrent chromosome VII region associated with the shared parent was excluded from the primary published-QTL comparison.

### Candidate-gene prioritization

Genes within QTL regions were identified using the S288C R64 annotation. Variants were annotated by coding consequence, PROVEAN score, and phyloP conservation.

## ACKNOWLEDGEMENTS

Thanks to all members of the Kruglyak Lab for helpful discussions and advice, and to Stephanie Moon and Ben Dodd. Thanks to Mark Arbing for the purified Tn5 and helpful discussions.

This work used computational and storage services associated with the Hoffman2 Cluster which is operated by the UCLA Office of Advanced Research Computing’s Research Technology Group.

Materials described in this publication (Tn5 enzyme) were prepared by the UCLA-DOE Protein Expression Technology Center which is supported by the U.S. Department of Energy, Office of Biological and Environmental Research (BER) program under Award Number DE-FC02-02ER63421.

## AUTHOR CONTRIBUTIONS

G.N.B.: Conceptualization; Data curation; Formal analysis; Funding acquisition; Investigation; Methodology; Project administration; Resources; Software; Supervision; Validation; Visualization; Writing – original draft; Writing – review & editing.

L.C.C.: Investigation; Methodology; Resources; Writing – review & editing. L.A.: Methodology.

H.M.: Data curation; Project administration; Resources. J.B.: Conceptualization.

J.S.B.: Conceptualization; Data curation; Formal analysis; Funding acquisition; Investigation; Methodology; Project administration; Resources; Software; Supervision; Validation; Visualization; Writing – original draft; Writing – review & editing.

L.K.: Conceptualization; Funding acquisition; Project administration; Resources; Writing – review & editing.

## FUNDING

L.K.: National Institutes of Health: 2RO1GM102308-06.

L.K. is a Howard Hughes Medical Institute Investigator

G.N.B. was a HHMI Hanna H. Gray Fellow in his postdoctoral phase for the duration of this work.

## DATA AND CODE AVAILABILITY

All results come from a single run of the analysis pipeline (v7), which records its configuration and writes every quality-control and peak table reported here.

BioProject ID: PRJNA1540837 https://www.ncbi.nlm.nih.gov/bioproject/1540837 Analyzed Data: 10.5281/zenodo.23193589

Code: https://github.com/brunig/microscale-xqtl Data Browser: https://brunig.github.io/microscale-xqtl/

